# Extracellular vesicles from Manila clam (*Ruditapes philippinarum*): tailored isolation from hemolymph and insights into water-derived vesicles

**DOI:** 10.64898/2026.05.29.727857

**Authors:** Valentina Moccia, Dalla Rovere Giulia, Thi Tran Ngoc Minh, Andrea Zendrini, Marije Kleinjan, Marc C. Roelofs, Paola Berto, Tzviya Zeev-Ben-Mordehai, Esther A. Zaal, Paolo Bergese, Annalisa Radeghieri, Massimo Milan, Marca H.M. Wauben, Valentina Zappulli

## Abstract

Extracellular vesicles (EVs) are evolutionarily conserved mediators of intercellular communication released by cells into biological fluids and the extracellular environment. Despite their growing relevance in biomedical and veterinary research, knowledge on EVs in marine bivalves remains limited. The aim of this study was to optimize tailored protocols for EV isolation from the hemolymph of the Manila clam (*Ruditapes philippinarum*) based on density gradient ultracentrifugation (dgUC) or size exclusion chromatography (SEC). EV-enriched fractions were identified through nanoparticle tracking analysis, protein quantification, transmission electron microscopy, and cryo-electron microscopy. Both methods successfully isolated small EVs (<200 nm). While dgUC yielded higher-purity preparations, SEC provided a higher recovery rate and compatibility with downstream metabolomic analyses. Metabolomics performed on SEC fractions and on hemolymph, revealed that EV-enriched fractions possessed a distinct metabolic signature including enrichment in metabolites associated with nucleotide metabolism, glycolysis, redox regulation, and energy metabolism. Furthermore, we performed a pilot investigation into the presence of EVs released into conditioned water by Manila clams. Using tangential flow filtration and ultrafiltration, EVs were successfully concentrated from water samples and characterized by nanoparticle tracking analysis, CONAN assay, atomic force microscopy, and electron microscopy. Our findings demonstrate the feasibility of isolating EVs both from Manila clam hemolymph and from conditioned water, providing the first evidence of water-derived EV recovery in aquatic animals. Although further methodological refinement is needed to improve the purity of EVs isolated from water, and additional characterization studies are required to better define the molecular composition of clam-derived EVs, these results establish a foundation for future investigations into the role of EVs in bivalve biology and their potential application as minimally invasive biomarkers for aquaculture, environmental monitoring, and ecosystem health assessment.

## Introduction

In the last two decades, extracellular vesicles (EVs) have emerged as complex intercellular shuttles conserved across organisms of different phyla and kingdoms^1–3^. EVs are membrane-enclosed nano-vesicles released by cells in the extracellular space^4–6^. By travelling through body fluids and interacting with close or distant “recipient” cells, EVs play a role in intercellular communication and participate in the regulation of both biological and pathological processes^5,6^. Beside their role in the regulation of physio-pathological processes, based also on their ability to carry part of the content of their cell of origin (e.g. proteins, RNAs, metabolites, lipids), one of the main applications of EVs is in liquid biopsy, where they are studied as promising diagnostic biomarkers^7–14^. To date, EVs have been detected in every biological fluid tested so far and have been demonstrated to be released by a variety of organisms including mammals, invertebrates, plants, fungi and bacteria^15–18^. Despite being known to be conserved mediators of intercellular communication, to date, most of the literature on EVs is related to applications in human medicine^16^. More recently, the studies of animal derived EVs is receiving increasing attention, however, the knowledge on EVs in certain animal species, such as invertebrates or wild animals, is still scarce ^15,16,18–20^.

Manila clams (*Ruditapes philippinarum)* are among the most farmed bivalves worldwide, having a consequent strong economic and social relevance, especially in coastal and riparian environments. The farming and trade of bivalves is of particular economic and social interest not only for their large distribution but also for their high nutritional profile and the low carbon footprint of the production chain^21,22^. These features make Manila clams, and more in general bivalves, cost-effective and sustainable foods^21,22^. In addition, bivalves have a filter feeding activity and resident behavior, which make them excellent candidates as environmental bioindicators for the monitoring of the effects of climate change or of environmental pollution ^23–25^. Despite the strong economic, social and environmental importance of bivalves, knowledge on extracellular vesicles in these organisms remains limited. Preliminary studies have described the presence and hypothetical roles of EVs in the hemolymph of some marine bivalve species ^26–29^. However, the hemolymph of marine bivalves is a complex biofluid, with distinct and different features compared to mammals’ which can make EV-purification challenging. Being present in all biofluids, EVs can also be released into the external environment during excretory activities. Once released, EVs retain molecular information on the releasing cell/organism, thereby representing valuable indicators of the physiological and pathological state ^30,31^. Given that bivalves possess an excretory apparatus, we can assume that they also release EVs into the surrounding aquatic environment and that these EVs may provide insights into their physiological or pathological conditions. Therefore, the analysis of EVs in seawater could represent a novel non-invasive approach to study bivalve biology and health status. Despite this promising potential, EV isolation from seawater has so far been reported only for marine bacteria derived-EVs, and the feasibility of isolating EVs released by aquatic animals has not been evaluated yet ^32–34^.

Considering the considerable translational potential of bivalve-derived EVs and the current lack of specific isolation methodologies, in this study we describe two tailored protocols for the purification of EVs from the hemolymph of Manila clams, one based on sucrose density gradient ultracentrifugation (dgUC) and one on size exclusion chromatography (SEC). To further investigate EV molecular composition in comparison with the whole hemolymph, the metabolomic profile of SEC-derived EVs was also analyzed. Moreover, given the limited knowledge regarding EVs released into the surrounding water by aquatic animals and the broad range of potential applications associated with their study, we additionally evaluated the feasibility of isolating EVs from conditioned water samples of Manila clams under laboratory conditions.

## Results

### Characterization of hemolymph derived extracellular vesicles

Hemolymph derived EVs were first purified by dgUC. Prior to dgUC, hemolymph was diluted using two different diluents,i.e. milliQ water supplemented with 3.2% of NaCl or PBS. The reason for comparing these two diluents is that during preliminary experiments, we noticed that the dilution of hemolymph in standard PBS caused the formation of (salt) precipitates. Here we assessed whether the use of PBS and the formation of precipitates during the initial phases of the EV isolation procedure hampered the recovery and purity of the EV sample.

To characterize hemolymph-derived nanoparticles, all individual fractions obtained by dgUC were collected. Based on previously reported EV-enriched fractions from other cellular sources isolated using comparable dgUC protocols^35^, fractions were pooled in groups of n=3 and subsequently characterized in terms of nanoparticle concentration, size distribution, and total protein content, with fractions 7–9 expected to represent the EV-enriched pool.

Using nanoparticle tracking analysis (NTA), pooled fractions from dgUC of hemolymph diluted in Milli-Q water supplemented with 3.2% NaCl or PBS (hereafter referred to as NaCl and PBS fractions) showed comparable results (Figure 1a, Table 1). The highest mean particle concentrations were observed in dgUC fractions 1–3 (high density fractions) for both PBS and NaCl (3.33×10^11^ ± 1.36×10^10^ and 2.31×10^11^ ± 2.35×10^10^ particles/ml, respectively). The remaining fractions displayed a progressive decrease in particle concentration (Table 1, Figure 1a). The mean particle size resulted comparable across all pooled dgUC fractions, ranging from 121.57 nm (PBS dgUC fractions 1–3) to 178.17 nm (NaCl dgUC fractions 4–6) (Table 1).

**Figure 1.**
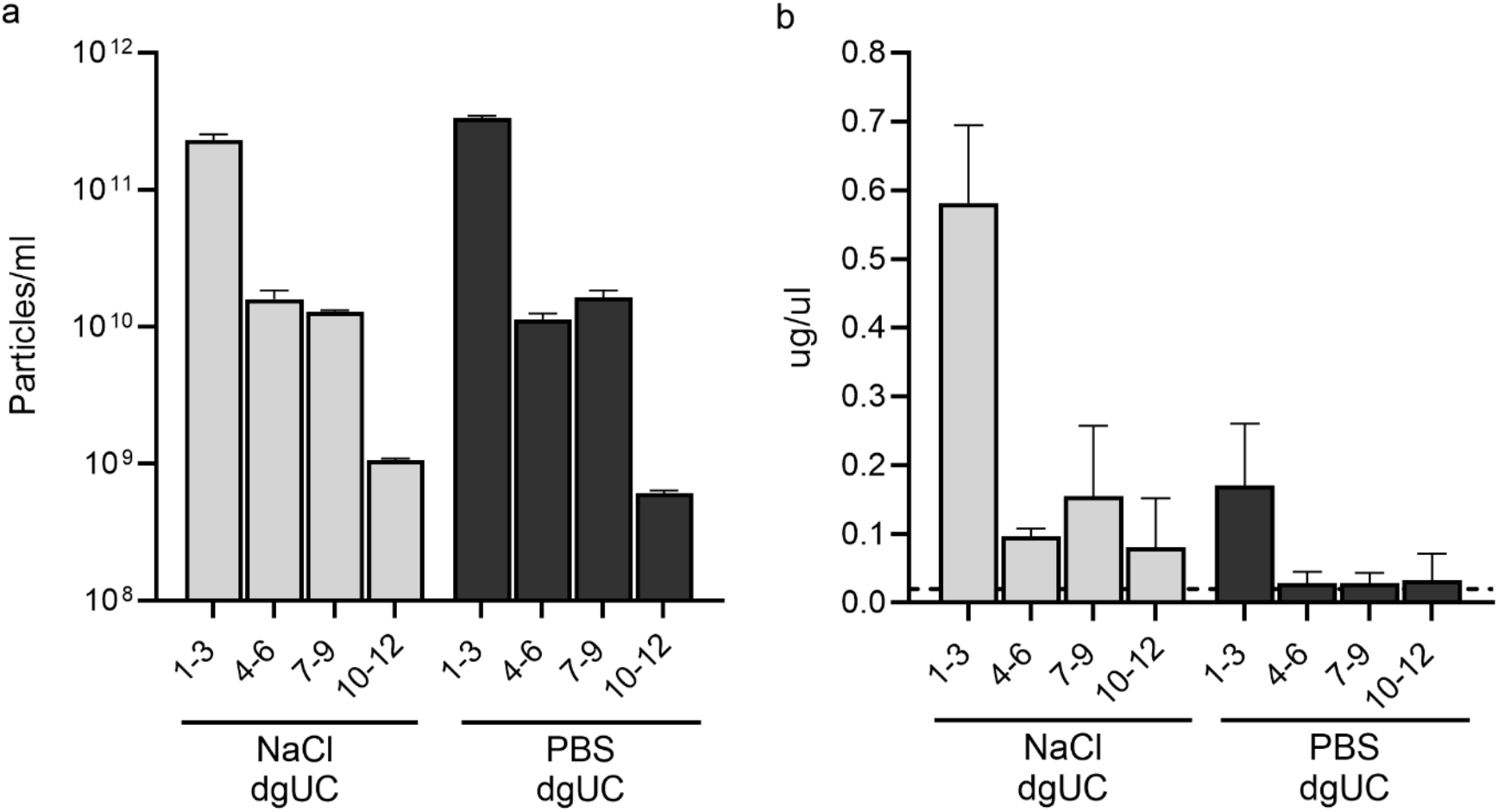
Histograms representing the mean particle (A) and protein concentration (B) in each pool of fractions (fr) derived from extracellular vesicle purification from Manila clam hemolymph with density gradient ultracentrifugation (dgUC) after hemolymph dilution in milliQ +3.2% of NaCl (NaCl dgUC) or in PBS (PBS dgUC). The highest particle and protein concentration was measured in both NaCl and PBS dgUC fr 1-3 and presented a decreasing trend in the later fractions.

**Table 1.** Results of EV characterization by Nanoparticle Tracking Analysis and protein quantification performed on pooled fractions of the EV isolation procedure from Manila clam hemolymph by density gradient ultracentrifugation (dgUC) or by size exclusion chromatography (SEC).

| Sample |  | Mean nanoparticle concentration (particle/ml) +/- SD | Mean nanoparticle size (nm) +/- SD | Mean protein concentration (ug/ul) +/- SD |
| --- | --- | --- | --- | --- |
| Hemolymph – NaCl dgUC fr | 1-3 | 2.31*10<sup>11</sup> +/- 2.35*10<sup>10</sup> | 124.17 +/- 9.23 | 0.58 +/- 0.11 |
|  | 4-6 | 1.59*10<sup>10</sup> +/- 2.52*10<sup>9</sup> | 178.17 +/- 1.03 | 0.10 +/- 0.01 |
|  | 7-9 | 1.29*10<sup>10</sup> +/- 2.74*10<sup>8</sup> | 149.57 +/- 1.87 | 0.15 +/- 0.1 |
|  | 10-12 | 1.06*10<sup>9</sup> +/- 2.76*10<sup>7</sup> | 163.43 +/- 2.33 | 0.08 +/- 0.07 |
| Hemolymph – PBS dgUC fr | 1-3 | 3.33*10<sup>11</sup> +/- 1.36*10<sup>10</sup> | 121.57 +/- 3.77 | 0.17 +/- 0.09 |
|  | 4-6 | 1.13*10<sup>10</sup> +/- 1.23*10<sup>9</sup> | 168.93 +/- 1.83 | 0.03 +/- 0.02 |
|  | 7-9 | 1.64*10<sup>10</sup> +/- 2.08*10<sup>9</sup> | 145.83 +/- 1.97 | 0.03 +/- 0.01 |
|  | 10-12 | 6.09*10<sup>8</sup> +/- 2.74*10<sup>7</sup> | 134.03 +/- 2.53 | 0.03 +/- 0.04 |
| Hemolymph - SEC fr | 1-5 | 9.35*10<sup>10</sup> +/- 3.01*10<sup>10</sup> | 139.4 +/- 1.93 | 0.51 +/- 0.08 |
|  | 6-10 | 4.51*10<sup>9</sup> +/- 2*10<sup>8</sup> | 129.3 +/- 3.57 | 0.08 +/- 0.04 |
|  | 11-15 | 8.18*10<sup>9</sup> +/- 2.8*10<sup>8</sup> | 132.73 +/- 3.2 | 0.11 +/- 0.02 |
|  | 16-20 | 1.72*10<sup>10</sup> +/- 1*10<sup>9</sup> | 151.63 +/- 2.67 | 0.11 +/- 0.06 |

SD: standard deviation, dgUC: density gradient ultracentrifugation, SEC: size exclusion chromatography, fr: pooled fractions

Protein quantification analysis revealed a trend similar to that observed with NTA, with dgUC fractions 1–3 showing the highest protein concentration (0.58 ± 0.11 µg/µl). In contrast to the overlapping measurements observed in NTA, protein concentrations in PBS dgUC fractions were generally lower than those in NaCl fractions (Table 1, Figure 1b).

To analyze EV-distribution across all the dgUC pooled fractions, negative staining transmission electron microscopy (TEM) was performed.

EVs were detected in all the dgUC pooled fractions but with differences in terms of purity. Both NaCl and PBS dgUC fractions 7-9 were confirmed as EV enriched fractions, as they contained a higher abundance of EVs and fewer co-isolated background particles compared to the other fractions (Figure 2). Across all conditions, EVs appeared as vesicular and rounded, cup-shaped structures predominantly with a size < 200 nm, consistent with the enrichment mainly of small EVs (Figure 2).

**Figure 2.**
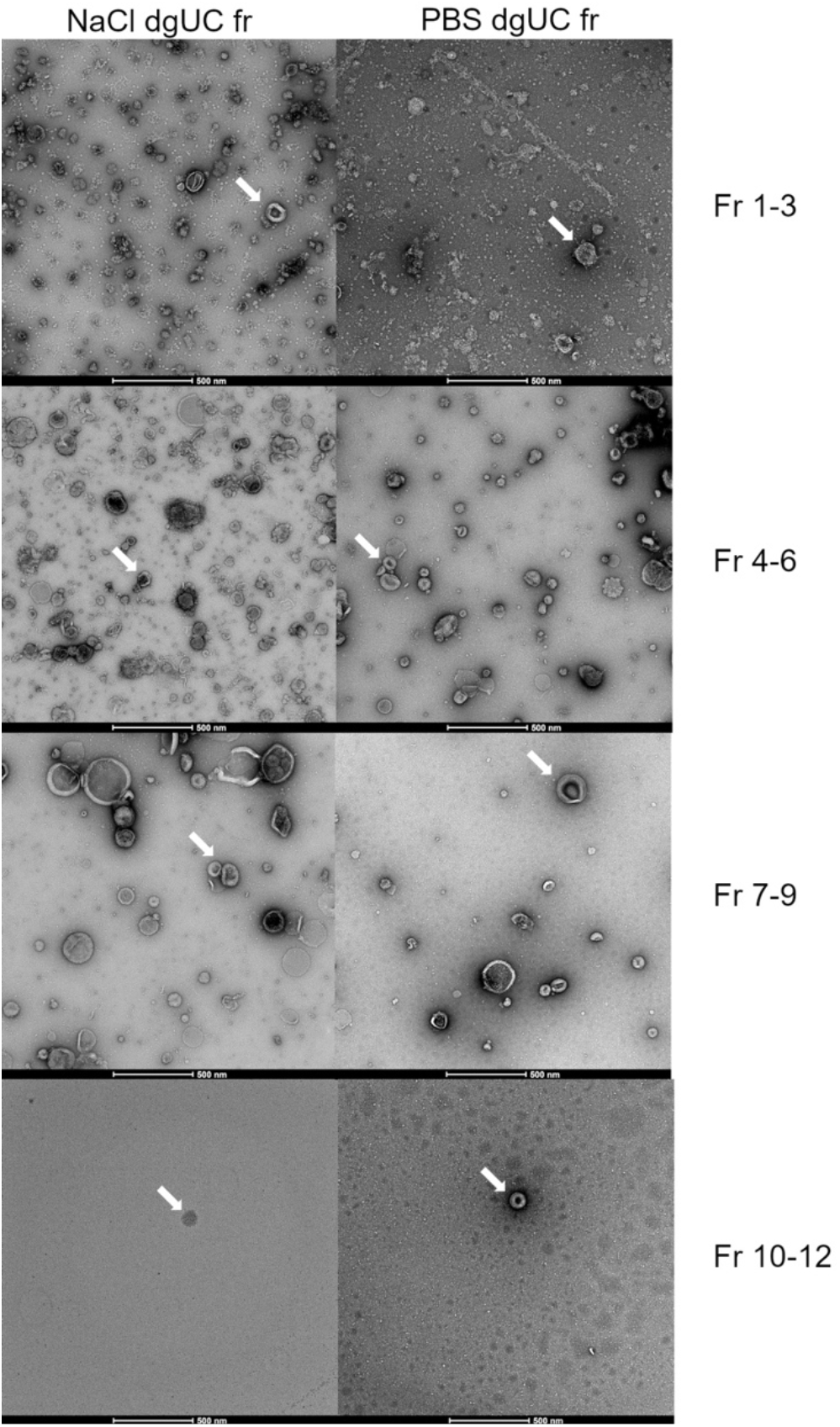
Electron microscopy (EM) performed on all density gradient pooled fractions (fr) derived from hemolymph diluted in milliQ+3.2% of NaCl (dgUC NaCl) or in PBS (dgUC PBS). Despite extracellular vesicles (EVs) were present across all fr, EVs with least co-purified contaminants in the background can be seen in fr 7-9 from both conditions. Arrows indicate examples of EVs in each fr.

**Figure 3.**
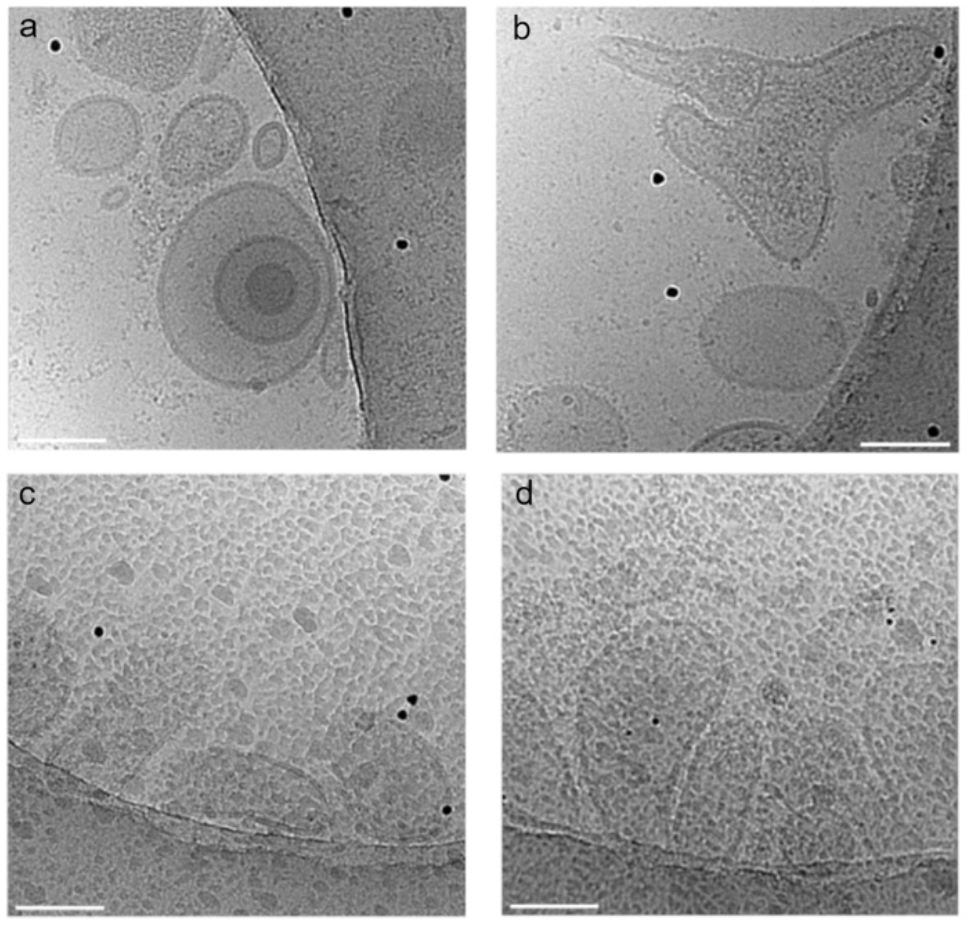
Cryo-EM of EVs from density gradient ultracentrifugation pooled fractions 7-9 of hemolymph diluted in milliQ+3.2% of NaCl (a, b), or diluted in PBS (c, d). EVs can be seen as vesicular structures of heterogenous shapes with an intact lipid bilayer. In PBS derived fractions, aggregates and precipitate-like structures on the surface hamper the quality of the figures (c, d). The scale bar is set at 100 nm.

**Figure 3.**
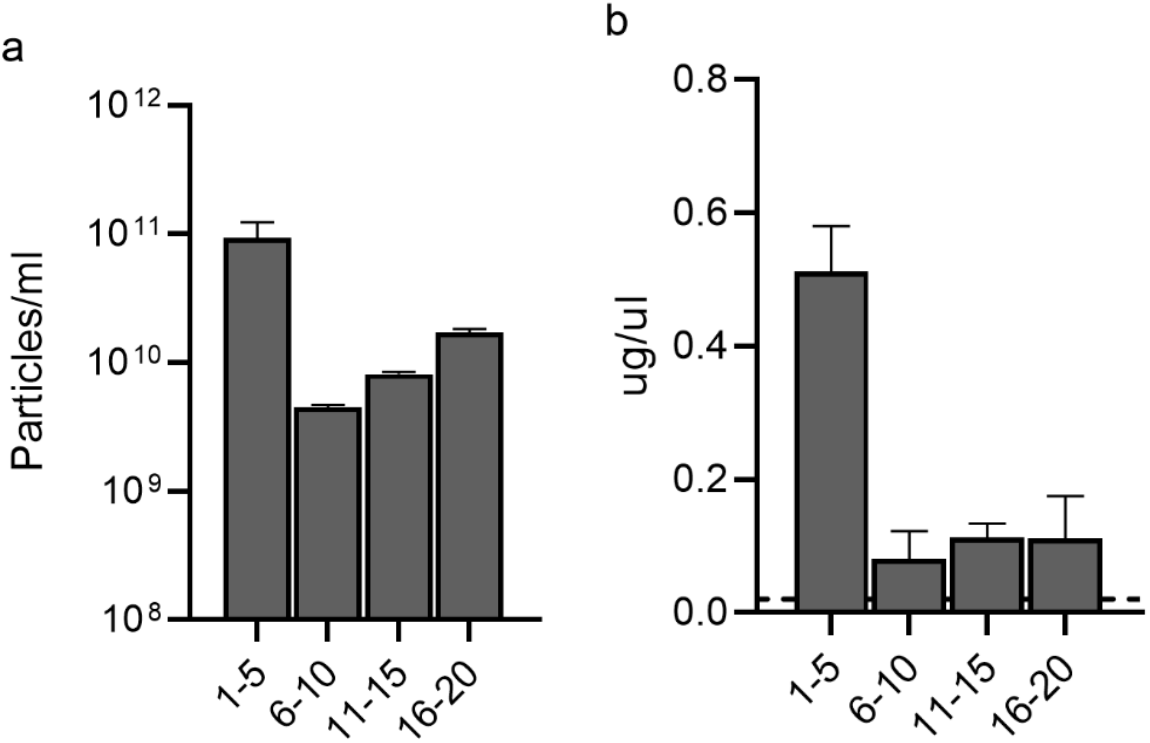
Histograms representing the mean particle (A) and protein concentration (B) in each pool of fractions (fr) derived from extracellular vesicle purification from Manila clam hemolymph with size exclusion chromatography (SEC). On the x axis, the numbers of the pooled SEC fr is present. The highest particle and protein concentration were measured in fr 1-5.

Following the identification of dgUC fractions 7-9 as EV-enriched fractions, Cryo-EM was performed to further assess potential differences in EV quality and integrity between NaCl and PBS dgUC fractions. Cryo-EM confirmed the preservation of EV integrity in both conditions, showing several EVs with an intact lipid bilayer (Figure 3). However, although EVs appeared structurally intact in both samples, PBS dgUC fractions 7-9 presented aggregates/precipitates in the background, which negatively affected image quality (Figure 3).

To further investigate EV-composition, we focused on metabolomic profiling, as metabolites represent one of the most conserved molecular classes and the Manila clam genome was not fully annotated yet at the time of the analysis. However, the presence of sucrose in dgUC fractions interfered with the quality of metabolomic analysis. Therefore, EVs were subsequently purified using a SEC-based protocol to enable the evaluation of the EV metabolite content.

For EV isolation using the SEC procedure, considering the results of Cryo-EM performed on NaCl and PBS dgUC fractions, hemolymph was diluted in milliQ water supplemented with 3.2% of NaCl. As performed for dgUC, to characterize nanoparticles in hemolymph, all individual fractions derived from SEC were collected and then pooled. Pools of n=5 fractions were made and characterized for nanoparticle concentration and size distribution, as well as total protein concentration, with fractions 1–5 expected to be EV-enriched pool based on the manufacturer’s specifications for the SEC column and on comparable protocols applied on other cellular sources^36^. At NTA, among SEC fractions, pool 1–5 exhibited the highest particle concentration (9.35×10^10^ ± 3.01×10^10^ particles/ml), with the following fractions 6-10 having the lowest particles concentration (4.51*10^9^ ± 2*10^8^, Table 1, Figure 3a).

Protein quantification analysis revealed a trend comparable to the NTA, with SEC fractions 1–5 presenting the highest protein level (0.51 ± 0.08 µg/µl) and fractions 6-10 the lowest (0.08 ± 0.04 µg/µl, Table 1, Figure 3b).

EV-distribution across SEC pooled fractions was then assessed by negative staining TEM which confirmed that most EVs with least contaminants were present in fractions 1-5 (Supplementary file).

Next, we aimed to verify the presence of canonical EV-associated proteins^37^. As markers, we selected CD9, Integrin-β, TSG101 and enolase. While enolase was selected due to its high conservation across species as a metabolic enzyme, CD9, Integrin-β and TSG101 are routinely used in our laboratory and have previously shown cross-reactivity with EVs purified from other animal species^38,39^. However, none of the tested antibodies showed cross-reactivity with either hemolymph-derived EVs or proteins extracted from clam tissues when analyzed by Western blotting (WB) (results not shown).

Then, to characterize EV associated molecules, metabolomics was performed. We determined the metabolic content of hemolymph derived-EVs and analyzed whether this content differed from the metabolic profile of hemolymph. LC-MS based metabolomics was then performed on pooled SEC fractions and on 10,000 *x* g hemolymph supernatant, since sucrose present in the dgUC fractions interfered with metabolite detection.

A comparable number of metabolites was detected across all samples, ranging from 62-81 metabolites (Figure 4a). The highest number of metabolites was detected in the EV-enriched SEC fractions, [SEC fractions 1-5 (81 metabolites)], followed by SEC fractions 15-20 (78 metabolites), hemolymph 10,000 *x* g supernatant (75 metabolites), SEC fractions 10-15 (69 metabolites) and SEC fractions 5-10 (62 metabolites) (Figure 4a).

**Figure 4.**
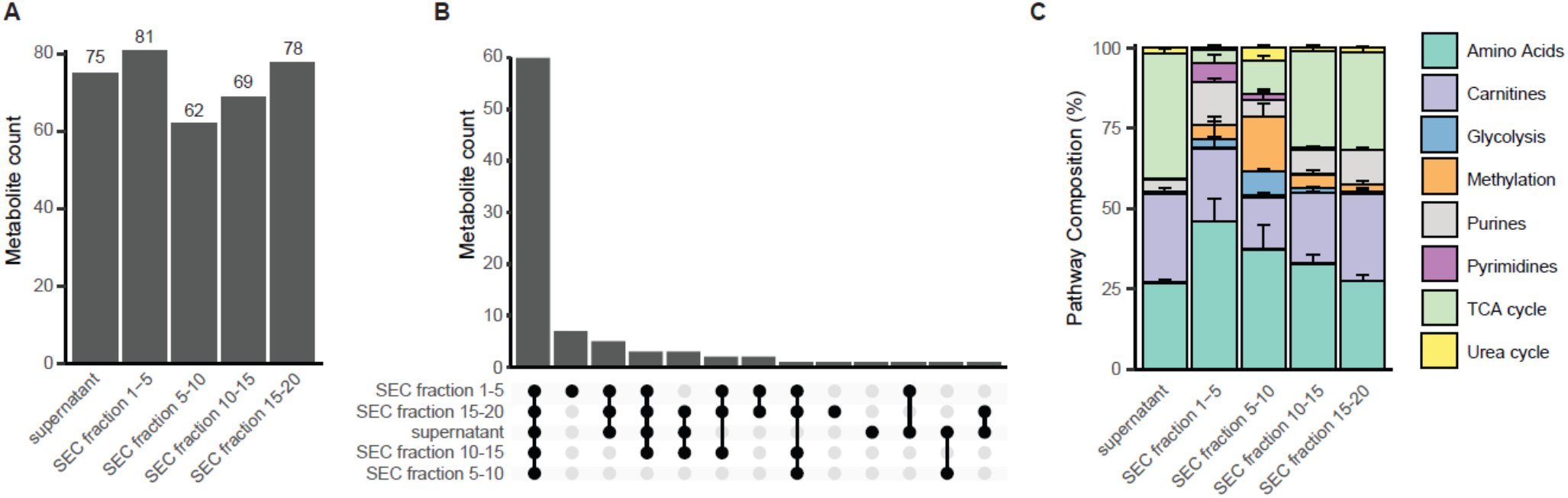
Metabolic profiling of hemolymph-derived extracellular vesicles (EVs) and size exclusion chromatography fractions. A) Total number of detected metabolites in pooled SEC fractions and the 10.000 x g hemolymph supernatant. B) UpSet plot showing the overlap of detected metabolites across SEC fractions and the hemolymph supernatant. Each bar represents the number of metabolites shared between the indicated sample combinations. C) Relative pathway composition of detected metabolites across samples. Metabolites were grouped into major metabolic pathways. Data represent pooled samples per fraction group.

The majority of the metabolites were shared between fractions. However, a subset of metabolites was uniquely detected in EV-enriched fractions (Figure 4b), i.e., UDP-N-Ac-Glucosamine, 2P-glycerate, glutathione, glucosamine-6P, NAD^+^, sedoheptulose7P, ADP, AMP (Figure 4b). These metabolites were not detected in the other fractions or in the hemolymph supernatant. To further characterize the differences in metabolic composition, detected metabolites were grouped by metabolic pathway (Figure 4c). While amino acid related metabolites constituted the largest proportion across all samples, the EV-enriched fractions showed a relative increase in metabolites associated with nucleotide metabolism, particularly purine and pyrimidine metabolites and glycolytic intermediates. In contrast, later fractions and the supernatant displayed a more balanced distribution across metabolic pathways, with more abundance of TCA cycle metabolites. Together, these data indicate that EV-enriched fractions exhibit a distinct metabolic composition as compared to the other SEC fractions and supernatant.

### Characterization of water released extracellular vesicles

To investigate the potential release of EVs into water by Manila clams, we performed a pilot study aimed at preliminary assessing both the presence of EVs in water and the feasibility of their isolation. Given the large volume of water (1L) compared with matrices commonly used for EV purification, an EV-concentration protocol based on tangential flow filtration (TFF) was applied. Subsequently, to further reduce the sample volume (from about 15 mL), samples were additionally concentrated by ultrafiltration. This protocol was applied to different water conditions: 1L containing 5 or 10 clams, Control 1 (C1) and Control 2 (C2). Controls were included to distinguish particles associated with the presence of alive clams from background particles potentially present in a non-completely sterile and closed water system (C1), and from particles potentially released from or associated with clam shells (C2). Each obtained sample was first characterized for nanoparticles size distribution at NTA and for EV-purity with CONAN assay ^40,41^. The highest particle concentration was detected in C2 (4.71*10^11^ +/-3.87*^11^ particles/ml), followed by EVs concentrated from 10 and 5 clam conditions (1.77 *10^11^ +/-2.29 *10^11^ and 1.47*10^11^ +/-1.43*10^11^ particle/ml, respectively) (Figure 5a). C1, presented the lowest particle concentration (2.42*10^10^ +/-1.82*10^9^ particle/ml) (Figure 5a) and was not included in the CONAN assay. C1 also presented an average particle size slightly higher compared to the other samples, with a detected mean size of 237 +/-6 nm compared to the other samples with similar size distribution (mean size in C2: 153 +/-4 nm; 5 clam sample: 180+/-3 nm; 10 clam sample: 160 +/-4 nm).

**Figure 5.**
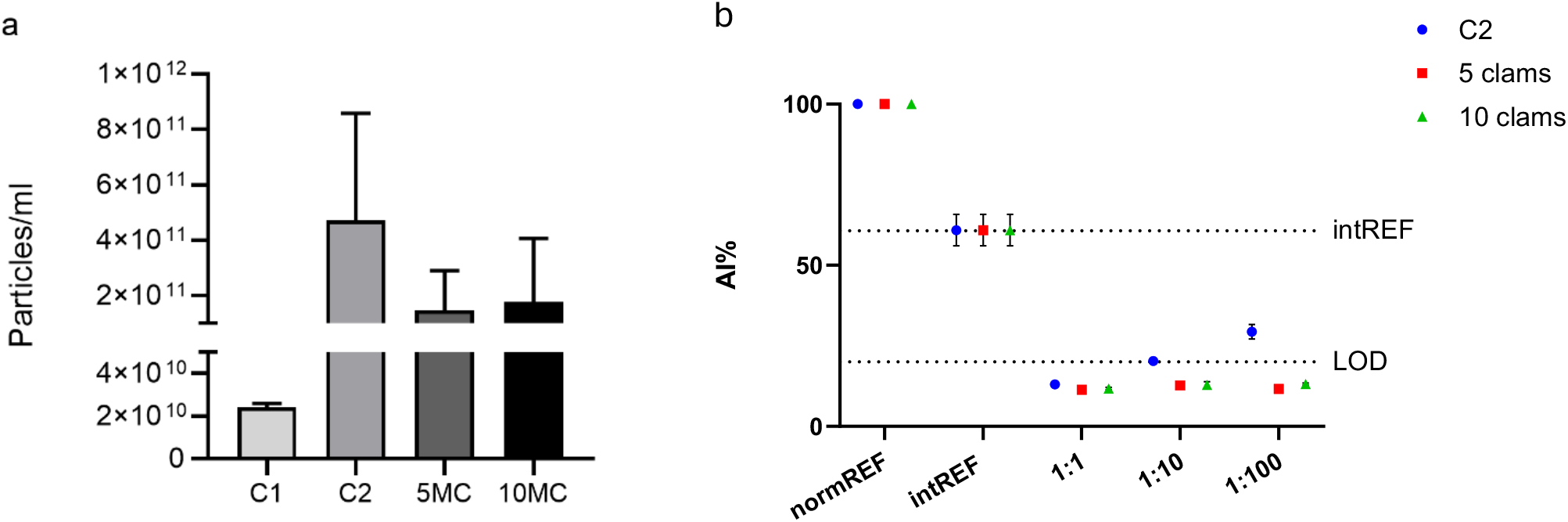
Characterization analysis performed on nanoparticles concentrated from water with 5 or 10 Manila clams (5MC, 10MC), from water with 10 empty shells (C2) and from only water samples (C1). Nanoparticle tracking analysis representing the mean concentration of nanoparticles/ml in each sample. The highest particle concentration was measured in C2, while the lowest in C1 (A). CONAN assay confirmed the absence of substantial amounts of soluble protein contaminants in all the three samples tested, suggesting C2 sample to be less concentrated in terms of membranous particles able to induce AuNP aggregation. AI = Aggregation index; NormaREF: pristine AuNP control; intREF: internal reference defining the level of aggregation of AuNPs induced solely by the buffer; LOD = Limit of detection defining the threshold for consider a sample pure from soluble proteins, in the limit of detection of the assay (B).

All the tested samples yielded positive results at the CONAN assay indicating that they were (i) sufficiently free of protein contaminants and (ii) contained particles capable of inducing citrate-capped gold nanospheres (AuNPs) clustering. Notably, the C2 sample (blue dots in Figure 5b) likely contained a lower particle concentration than the samples derived from 5 and 10 clams (red squares and green triangles respectively in Figure 5b), as a higher sample concentration was required to induce AuNP aggregation.

The presence of EV-like structures in samples derived from 5 and 10 clams was confirmed by in air-AFM (Figure 6, white arrows). On the other hand, in air-AFM showed that C2 sample contained mostly aggregated material not ascribable to EVs (Figure 6, red arrows), with a very minor quantity of objects carrying EV-like features (Figure 6, white arrows), results in line with those obtained from the CONAN assay.

**Figure 6.**
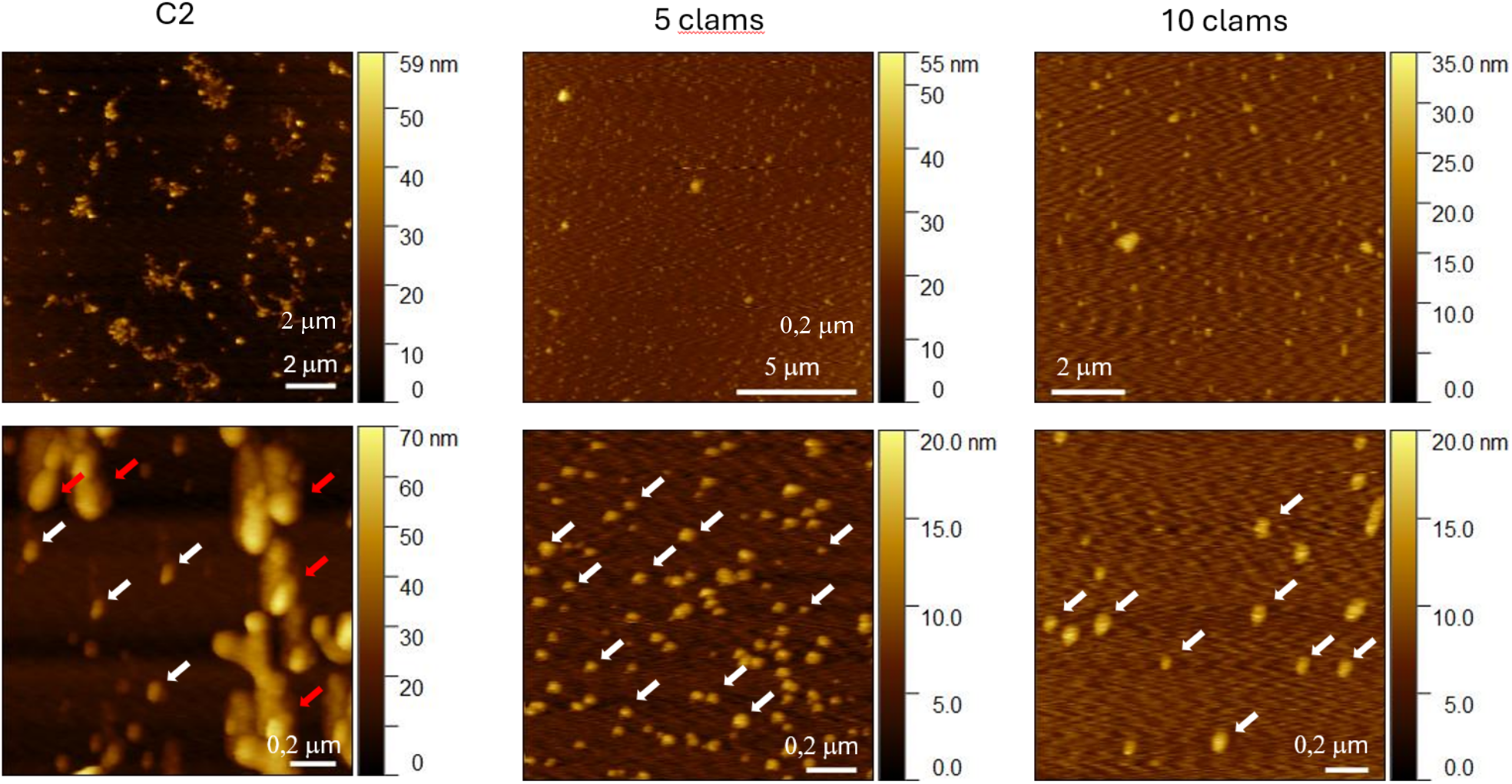
Representative in-air Atomic Force Microscopy micrographs of particles concentrated from water with empty shells (C2), with 5 clams or 10 Manila clam samples. Extracellular vesicle-like structures are indicated by white arrows; red arrows indicate aggregates of unknown origin.

Finally, to visualize EV presence and the nature of potential co-isolated particles, all water derived samples were analyzed by TEM with negative staining. EVs were detected in all samples but appeared more frequently in comparison with co-isolated material in the 5 and 10 clam sample (Figure 7a-d). In all conditions, additional variably shaped heterogeneous nanostructures were present (Figure 7a-d). Of note, several whitish elongated acicular (100-200 nm) structures were detected in C2 but were absent from the other samples (Figure 7b). In the 5 and 10 clam samples, biological nanostructures, probably consistent with proteins and viral structures, were also present (Figure 7c-d).

**Figure 7.**
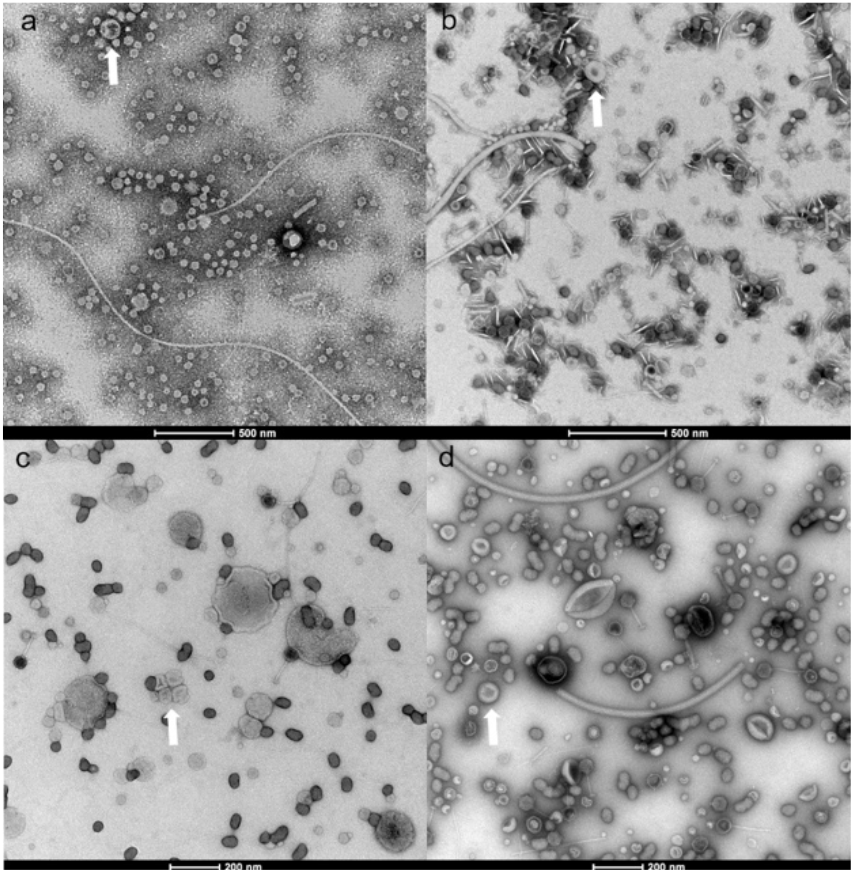
Negative staining transmission electron microscopy performed on particles concentrated from only water (sample C1, a), from water containing 10 empty Manila clam shells (sample C2, b) or from water containing 5 (c) or 10 Manila clams (d). White arrows indicate extracellular vesicles (EVs). In all conditions, other heterogenous nanostructures co-isolated with EVs.

## Discussion

In this study we describe two effective tailored protocols for EV isolation from Manila clam hemolymph and provide the first insights into the metabolite profile of clam derived-EVs. In addition, we preliminary investigated the presence of EVs in conditioned water with clams under controlled laboratory conditions and evaluated the feasibility of their isolation.To date, EVs have received considerable attention in human medicine and in several animal species; however, they have been only scarcely explored in bivalves and have never been investigated in Manila clams^16^. Some studies have analyzed EVs in the hemolymph of other marine bivalve species, but purification has mainly relied on ultracentrifugation or exosome purification kits designed for serum. The resulting EV preparations were not thoroughly characterized, particularly for the presence of possible co-isolated contaminants^28,29,42,43^. In the present study, we successfully purified EVs from Manila clam hemolymph using protocols based on dgUC or SEC. Given the absence of literature describing these two methodologies applied to hemolymph, we characterized all dgUC and SEC fractions to assess EV distribution. In addition, given that in preliminary experiments performed in our laboratory the dilution of hemolymph in PBS caused the formation of salt precipitates, we also evaluated the effects of diluting hemolymph in PBS or in an isosmotic buffer with Manila clam biofluids (milliQ supplemented with 3.2% of NaCl), to determine possible differences in terms of isolated EV quantity and quality.

dgUC was the first protocol applied to EV purification from hemolymph. NTA, protein quantification analysis and TEM performed across all dgUC pooled fractions yielded consistent results and allowed us to identify dgUC fractions 7-9 as EV enriched fractions among both NaCl and PBS dgUC pooled fractions^35,44^. TEM particularly confirmed that fractions 7–9 contained EVs with higher purity despite lower particle and protein concentrations compared with fractions 1–3. These findings are consistent with the working principle of dgUC and with observations reported for EVs isolated from other biological matrices, where denser proteins and contaminants accumulate in the lower fractions of the gradient ^35,44^. The results further suggest that Manila clam-derived EVs exhibit densities comparable to those reported for mammalian EVs. Although NTA, protein quantification, and TEM analyses did not reveal major differences between PBS and NaCl dilution conditions, Cryo-EM demonstrated the presence of salt-like precipitates and aggregate structures in PBS-derived fractions 7-9, which negatively affected image quality. While previous studies on marine bivalve EVs have commonly employed PBS without reporting similar issues, our observations indicate that the use of isosmotic buffers may be preferable when processing biofluids from marine species like bivalves, as precipitation artifacts may interfere with downstream analyses. Similar improvements associated with osmolarity-adjusted reagents have also recently been reported for EV isolation from freshwater fish biofluids ^45^.

Because a fully annotated Manila clam genome was not available at the time of the analysis, metabolomics was selected as an alternative strategy to characterize EV cargo, given the high degree of metabolite conservation across species. However, the presence of sucrose in dgUC fractions interfered with metabolomic analyses, prompting the development of a SEC-based protocol for EV purification. NTA, protein quantification analysis and TEM were performed also across all SEC pooled fractions and allowed us to identify the EV enriched fractions in SEC fractions 1-5^36,44,46,47^. SEC fractions 1-5 displayed the highest particle and protein concentration and TEM confirmed the presence of most EVs. In contrast, later fractions contained fewer EVs and were enriched in soluble proteins and aggregates. These observations are consistent with the established principles of SEC, whereby smaller molecules and proteins elute later, whereas larger vesicular particles elute earlier ^36,44,46–49^. Compared with dgUC, SEC yielded higher particle and protein concentrations in EV-enriched fractions, although TEM analysis indicated that dgUC provided cleaner EV preparations with reduced background contamination. This observation aligns with previous reports showing that dgUC generally produces EV preparations of higher purity, albeit often at the expense of lower vesicle recovery^37^.

Notably, no cross-reactivity was observed between Manila clam proteins and some commonly used mammalian EV markers, including Integrin-β, CD9, TSG101, and enolase. Previous studies have reported the detection of CD9 and CD63 in EVs isolated from the hemolymph of other bivalve species (*Pinctada fucata martensii, Mytilus edulis, Mya arenaria, Crassostrea virginica, Ensis leei)*^29,50^. Although EV proteins are highly conserved among mammals, bivalves are phylogenetically distant from humans, which are the target species for most commercial antibodies, and the sequence of homology between human and bivalve proteins is generally low according to protein databases (e.g. Uniprot). However, further BLAST analysis specifically comparing bivalve EV protein sequences with antibody host targeted species are needed to better identify cross-reactivity between bivalve EV proteins and commercial antibodies. Given the challenges associated with protein-based characterization, metabolomic analyses were performed on SEC fractions and on hemolymph supernatant to investigate differences between EV-associated and non-EV-associated metabolites. EV-enriched SEC fractions 1-5 contained the highest number of detected metabolites and included a subset of metabolites not observed in other fractions. This supports the notion that EV-associated fractions possess a distinct metabolite composition as compared to the surrounding hemolymph. Notably, several of the uniquely detected metabolites (NAD+, AMP, Glutathione) are central to cellular energy metabolism and redox regulation, suggesting a potential role for EVs in transferring metabolites involved in metabolic signaling^51^. Although soluble metabolites may co-elute with EVs during SEC separation or may leak out of EVs, the observed differences in composition between EV-enriched fractions and later fractions or supernatant indicate selective enrichment of specific metabolites in EV-associated fractions. Compared with other omics approaches such as proteomics or RNA sequencing, metabolomics has been less frequently applied to the study of EV cargo.

Nevertheless, several studies in human medicine have investigated EV-associated metabolites, particularly in the context of identifying cancer diagnostic biomarkers^14,52,53^. These studies have demonstrated that EV metabolite cargo can reflect different disease state and can vary under different physiological conditions, such as after intense physical exercise^54^. The classes of metabolites detected in SEC EV-enriched fractions in our study, including amino acids, nucleotide components, carnitines and metabolites involved in glycolysis, methylation, TCA and urea cycles, have also been described in human EVs derived from various sources, including cell cultures, plasma and urine^14,52–54^. Notably, carnitines, amino acids and glycolysis related metabolites have also been detected in EVs isolated from macaque plasma and from *Bacterioides* culture medium, suggesting that EVs might play conserved roles in these metabolic pathways across different species^14^.

Because organisms can release EVs into the environment through excretory processes, we also preliminarily investigated the presence of EVs into water conditioned with Manila clams and assessed the feasibility of their isolation. Both seawater and clams constitute highly complex biological systems that may contain bacteria, viruses, algae, and other microorganisms capable of releasing extracellular particles^17,32,55–57^. To account for this complexity, two control conditions were included to mimic nanoparticles naturally present in non-sterile aquatic systems (C1) and particles potentially originating from organisms associated with clam shells (C2). In addition, two different clam densities (5 or 10 clams in 1L) were tested to evaluate if EV-shedding was influenced by animal density. NTA detected a higher particle concentration in C2 compared with the samples containing live clams, whereas no substantial differences were observed between the 5-clam and 10-clam conditions. The absence of clear density-dependent differences may be related to the relatively short incubation time or to possible re-filtration of EVs by the clams themselves. Importantly, CONAN assay and AFM analyses demonstrated that most particles detected in C2 were not membranous structures compatible with EVs, whereas EV-like membranous particles were observed in samples containing live clams. TEM further revealed elongated acicular structures in C2, likely contributing to NTA particle counts. Based on their morphology and on their absence in the other conditions, these structures may be mineral particles originating from empty clam shells, which are primarily composed of calcium carbonate and organic matrix components^58^. Furthermore, because control samples did not contain live clams, EVs detected in controls may have originated from microorganisms associated with shells or with the experimental system. TEM analyses additionally revealed the presence of viral particles and other unidentified nanoparticles across all conditions. Further studies will therefore be required to clarify the origin and composition of these particles and to optimize EV purification protocols from aquatic environments. To our knowledge, this study represents the first attempt to purify EVs released into water by animals. Previous studies have primarily focused on the purification of bacterial EVs from large volumes of ocean water to investigate their genetic content or to assess how different environmental conditions can modulate their release and composition^32–34^. Although substantial methodological improvements are still required to separate EVs from non-EV nanoparticles and to identify their cellular origin, our findings demonstrate that clams, together with associated microorganisms, release EVs into the surrounding water and that their isolation is feasible even from relatively small water volumes under laboratory conditions.

In conclusion, we describe two tailored protocols for EV purification from Manila clam hemolymph and provide preliminary evidence supporting the feasibility of EV isolation from seawater conditioned with clams. Our results suggest that the use of isosmotic buffers may improve EV sample quality when working with marine bivalves. Moreover, we demonstrate that metabolomic profiling represents a valuable alternative strategy for EV characterization in species lacking fully annotated genomes or proteomes. The recent availability of improved genomic and proteomic resources for *Ruditapes philippinarum* will now facilitate more comprehensive investigations of EV-associated protein markers, RNAs, and microRNAs, enabling deeper understanding of EV-mediated regulatory mechanisms in bivalves. Finally, although additional studies are required to fully characterize EVs and non-EV particles present in aquatic environments, the analysis of water-derived EVs represents a promising non-invasive approach for investigating aquatic organisms and ecosystems.

Further improvements of such approaches may have important applications in environmental monitoring, conservation biology, aquaculture, and the study of organismal responses to environmental stressors, including climate change, heat waves, and emerging contaminants.

## Methods

### Hemolymph collection and extracellular vesicle isolation

Fresh hemolymph was extracted from commercial adult alive Manila clams *(Ruditapes philippinarum)* using a non-lethal method. Briefly, a 1mL syringe attached to a 25-gauge needle was used to collect hemolymph from the posterior adductor muscle by inserting the needle through the hinge ligament edge, which was distinguished from other tissues by the given resistance at the withdrawal^59^. 200uL – 1mL of hemolymph were extracted from each Manila clam and pooled in aliquots of 4.5 mL collected from nine clams. Hemolymph aliquots were then centrifuged at 800 *x g* for 10 minutes to discard hemocytes and the supernatant collected and stocked at −80°C until EV purification by dgUC or by SEC.

For both EV isolation through dgUC or SEC, thawed hemolymph samples were centrifuged at 2,000 *x* g for 10 minutes at 4°C to discard large cell debris. The pellet was discarded and the supernatant transferred in ultracentrifuge tubes (Beckman Coulter) and ultracentrifuged at 10,000 *x* g for 30 minutes at 4°C (Optima L-90 K, Beckman Coulter) in a swinging bucket rotor (SW60ti, Beckman Coulter). Then, for EV purification with dgUC, 3.25 ml of the 10,000 *x* g supernatant was diluted 1:1 with either milliQ water supplemented with 3.2% of NaCl or with PBS both double filtered with a 200nm filter, in order to compare the use of a diluent isosmotic with hemolymph with standard PBS. The solution was loaded on the top of a sucrose density gradient, which was made by layering successive sucrose solutions of decreasing density (2.0 M – 0.4 M) on top of 2.5 M sucrose. The gradient was then ultracentrifuged at 200,000 *x* g for 16 hours at 4°C (Optima L-90 K, Beckman Coulter) in a swinging bucket rotor (SW40ti, Beckman Coulter). Density fractions were collected and three fractions of 500 uL pooled together for both dilution conditions as follows: high-density fractions 1–3 (mean density: 1.28– 1.26 g/mL), fractions 4–6 (mean density: 1.24–1.20 g/mL), EV-enriched fractions 7–9 (mean density: 1.18–1.13 g/mL), and low-density fractions 10–12 (mean density: 1.12–1.08 g/mL). The pooled density gradient fractions were diluted in 11.5 mL of dfPBS, transferred to ultracentrifuge tubes (Beckman Coulter) and ultracentrifuged at 100,000 × *g* for 65 min at 4°C (SW40ti rotor, Beckman Coulter, Optima L-90 K, Beckman Coulter). The obtained pellets were then used for further experimentation.

To perform EV-isolation with SEC, 3.25 mL of the 10,000 *x* g supernatant were concentrated with 100 kDa ultrafiltration tubes (Amicon Ultra centrifugal filters, Merck Millipore, Burlington, MA, USA) by centrifugation at 3,000 x g for 20 minutes to reach a volume of 500 uL and then loaded on the top of qEVoriginal/70 nm columns (IZON Science). SEC was performed according to manufacturer’s instructions and 400 ul fractions collected and pooled as follows: 1-5; 6-10; 11-15; 16-20. The pooled SEC fractions were then diluted in 11 mL of PBS, transferred to ultracentrifuge tubes (Beckman Coulter) and ultracentrifuged at 100,000 × *g* for 65 min at 4°C (SW40ti rotor, Beckman Coulter, Optima L-90 K, Beckman Coulter) to collect the pellets which were used for further analysis.

### Experimental set-up and extracellular vesicle concentration from water

To concentrate EVs released from Manila clams in water, a laboratory experimental set-up was created. Manila clams with a similar size (ranging from 5 to 7 grams per clam) were grouped in two different concentrations, 5 or 10 clams, and kept at room temperature (21°C) in a sterilized closed bottle containing 1L of deionized water added with 3.2% of NaCl and an oxygenator. To compare particles released from alive clams to the background particles normally present in water and to those released from organisms naturally co-existing with clam shells (e.g. bacteria, algae), two different control conditions were included: a. Control 1 (C1) consisting of 1L of deionized water added with 3.2% of NaCl and an oxygenator; b.Control 2 (C2), consisting of 1L of deionized water added with 3.2% of NaCl, an oxygenator and 10 Manila clam empty shells. Shells were previously mechanically cleaned with brushes between the valves to remove possible tissue fragments. After 48 hours, clams were removed from the bottles and water processed for EV-concentration. Each sample was first filtered with a 0.45um filter, then concentrated ~ 100 times by Tangential Flow Filtration (TFF) (performed at the dept. of Molecular and Translational Medicine, University of Brescia, Italy) using a Repligen KrosFlo system equipped with a MicroKros® 500 kDa mPES filtering cartridge. Cartridge integrity was verified according to the manufacturer’s instructions, and the system was flushed with 2.5 L of sterile deionized water prior to use, at a flow rate of 100 mL/min in manual mode. Sample concentration was carried out in concentration mode (C mode), with a transmembrane pressure (TMP) maintained between 0.4 and 0.5 and a flow rate ranging from 100 to 135 mL/min, with automated back-pressure valve control enabled. After concentration of each sample, the cartridge was cleaned with 0.2 M NaOH (manual mode; 500 mL flushed to waste followed by recirculation for 15 min at a flow rate of 100 mL/min). The system was then rinsed with sterile PBS until pH neutralization was achieved, followed by a final rinse with 1 L of sterile water before processing the next sample.

The resulting retentate (approximately 15 ml from 1 L) was further concentrated by UF using 100 kDa filters. The obtained sample was then further analyzed by NTA, CONAN Assay, negative staining TEM and in Air-AFM.

### Nanoparticle Tracking Analysis and protein analysis

After EV-isolation, all the hemolymph pellets from the dgUC or SEC pooled fractions and EVs concentrated from all water samples were characterized for particle concentration and size distribution by NTA.

The pellets were dissolved in 100 ul of PBS and analyzed for particle concentration and size distribution with NanoSight NS300 (Malvern). Samples were progressively diluted in PBS to reach the correct dilution to gain reliable measurements. For each sample, camera level was set at 12, and three movies of 60s each were recorded and analyzed using the 3.4 NTA software. For particle quantification, the detection threshold was set at 5 and measurements were considered reliable when within the following instrument optimal working ranges: particles per frame from 20 to 120; particle concentration between 10^6^ and 10^9^ per mL; ratio of valid particles to total particles higher or equal to 1/5.

All hemolymph derived dgUC and SEC fractions were also analyzed for protein concentration. To perform the analysis, each pellet was dissolved in 60 ul of RIPA buffer (ThermoFisher Scientific) supplemented with protease inhibitor and protein concentrations were calculated using the Pierce BCA protein Assay Kit (ThermoFisher Scientific) according to manufacturer’s protocol.

The purity of the EV-samples concentrated from water, with respect to soluble protein contaminants, was assessed using the CONAN assay (Department of Molecular and Translational Medicine, University of Brescia, Brescia, Italy) following the protocols described by Zendrini and coauthors^41^. Briefly, the assay exploits the spectroscopic properties of AuNPs to assess the presence of membranous particles in solution, their relative abundance, and their purity with respect to soluble protein contaminants. In pure EV preparations, AuNPs adsorb to and cluster at the EV membrane. In contrast, when exogenous protein contaminants are present, AuNPs are preferentially coated by these species, which inhibits nanoparticle clustering. AuNP aggregation induces a red shift in the localized surface plasmon resonance (LSPR) absorption peak, resulting in a colour change of the solution from red to blue that is proportional to the purity of the EV preparation. AuNP clustering can be quantified using the Aggregation Index (AI), which directly correlates AuNP aggregation with preparation purity. The AI is calculated from UV/Vis spectra and is defined here as the ratio between the absorbance at 519 nm and the sum of the absorbances at 650 nm and 850 nm (AI = A_519_ / (A_650_+ A_850_)). The AI decreases as the solution colour shifts from red to blue and is inversely proportional to sample purity. When the preparation is sufficiently pure, the AI can also provide information on the relative abundance of membrane material in solution ^40,41^.

NTA and protein quantification analysis performed on hemolymph derived samples were performed in three biological replicates per each dgUC and SEC pool of fractions, whereas NTA and CONAN assay were performed on water derived samples in biological duplicates.

### Electron Microscopy

Negative staining TEM was performed on all pooled dgUC and SEC fractions for hemolymph derived samples and on each EV sample from the different water conditions to assess EV-presence and distribution.

TEM was performed in different institutions. Hemolymph derived dgUC fractions were analyzed at the Bijvoet Center for Biomolecular Research (Utrecht, the Netherlands), hemolymph derived SEC fractions at the Istituto Zooprofilattico delle Venezie (Legnaro (PD), Italy) while water derived samples were analyzed in service at the department of Biology of the University of Padova (Padova, Italy).

Each EV-enriched pellet was dissolved in 20 ul of milliQ added with 130 mM NaCl and 25 mM of HEPES (pH 7.4) on ice. Subsequently for TEM performed on dgUC fractions and water derived samples, 4 μl of sample was applied to carbon-coated copper grids that were glow discharged (PELCO easiGlow, Ted Pella). The sample was allowed to absorb for 30 seconds prior to 2x wash with MQ and 3x staining with 2% uranyl acetate solution, with the last washing step lasting for ~30 seconds. Grids were allowed to dry in air for ~5 min and then imaged on a FEI Talos L120C transmission EM operated at 120 keV, equipped with a CCD camera.

TEM analysis on SEC fractions was performed using a Fei Tecnai 12 Spirit Instrument (FEI Company-Hillsboro OR,USA). Briefly, 20 µl of EV-enriched fractions were centrifuged for 5 min with a 200-mesh copper grid with formvar and carbon coatings (Electron Microscopy Sciences, Hatfield, PA, USA). After blotting the liquid in excess, sample was subjected to negative staining for 1 minute with a 2% solution of sodium phosphotungstate that was removed immediately after. The grid was subsequently dried for 5 min and examined at 120 keV.

To assess EV-integrity, Cryo-EM was performed on hemolymph dgUC EV fractions 7-9 (Bijvoet Center for Biomolecular Research, Utrecht, the Netherlands). The samples were diluted as previously described for negative staining TEM. 4 µL of sample was loaded onto a Quantifoil 2/1 300 mesh grid (Quantifoil Micro Tools) that was glow discharged (PELCO easiGlow, Ted Pella). 1 µL of a BSA-conjugated 10 nm gold beads (Aurion) suspension was added and the drop was blotted from the back (other side of sample deposition) for 4-6 seconds. Sample was vitrified by plunge freezing in liquid ethane-propane mix (37% ethane) using a manual plunge-freezer (MPI-Martinsried). Data was collected on a Talos Arctica (Thermo Fisher Scientific) transmission electron microscope operated at 200 kV, equipped with a postcolumn energy filter (Gatan) operated in zero-loss imaging mode with a 20-eV energy selecting slit. Projection images were collected using a K2 Summit direct electron detector (Gatan) in counting mode with dose fractionation, at a magnification of 100,000x (1.359 Å/pix). Target defocus was set at −4 micrometers and total dose approximately 50 e-/Å.

In-air AFM imaging was performed on water derived samples using a Nanosurf NaioAFM equipped with a Multi75-AI-G tip (Budget Sensors) (Department of Molecular and Translational Medicine, University of Brescia, Brescia, Italy). Samples were centrifuge at 100k x *g* for 90 minutes. Pellets were then resuspended in 100 uL and diluted 10 to 50 times sterile water μL (Milli-Q, Merck Millipore). Aliquots of 5 μL were deposited onto freshly cleaved mica sheets (Grade V-1; thickness 0.15 mm; size 15 × 15 mm^2^) and dried at RT for 30 minutes. Images were acquired in tapping mode, with scan sizes ranging from 1.5 to 25 μm and a scan speed of between 0.8 and 1.2 s per scanning line. Instrument sensitivity was set to 55%, and P-, I-, and D-gain to 5000, 500 and 0, respectively. Image processing was performed using Gwyddion (version 2.61).

### Metabolomics

To investigate the composition of hemolymph derived-EVs, metabolomics was performed at the dept. of Biomolecular Health Sciences (Utrecht University, Utrecht, The Netherlands). Since the presence of sucrose in dgUC fractions hampers the quality of metabolomic analysis, metabolomics was performed on hemolymph pooled fractions isolated by SEC and on the 10,000 *x* g supernatant, to assess the difference in metabolite composition in the different SEC fractions, in EVs and in the rest of hemolymph. The pellet derived from each SEC pool of fractions was dissolved in 10 ul PBS and lysed by the addition of 40 ul methanol/acetonitrile (1:1). For hemolymph supernatant, 100 ul were used and processed in the same way, by adding 100 ul methanol/acetonitrile (1:1). Samples were then centrifuged at 16.000 *x* g for 15 minutes at 4 °C to remove cell debris and proteins and supernatants were collected for LC-MS analysis.

LC-MS analysis was performed on a Q-Exactive HF mass spectrometer (Thermo Scientific) coupled to a Vanquish autosampler and pump (Thermo Scientific). The MS operated in polarity-switching mode with spray voltages of 4.5 kV and −3.5 kV. Metabolites were separated using a Sequant ZIC-pHILIC column (2.1 x 150 mm, 5 μm, guard column 2.1 x 20 mm, 5 μm; Merck) with elution buffers acetonitrile (A) and eluent B (20 mM (NH4)2CO3, 0.1% NH4OH in ULC/MS grade water (Biosolve)). Gradient ran from 20% eluent B to 60% eluent B in 20 minutes, followed by a wash step at 80% and equilibration at 20%. Flow rate was set at 100 μl/min. Analysis was performed using Tracefinder software (Thermo Scientific). Metabolites were identified and quantified on the basis of exact mass within 5 ppm and further validated by concordance with retention times of standards. Peak intensities were normalized based on total ion count.

## Notes

### Competing Interest Statement

The authors have declared no competing interest.

